# Mechanistic modeling of the SARS-CoV-2 and immune system interplay unravels design principles for diverse clinicopathological outcomes

**DOI:** 10.1101/2020.05.16.097238

**Authors:** Sarthak Sahoo, Kishore Hari, Siddharth Jhunjhunwala, Mohit Kumar Jolly

## Abstract

The disease caused by SARS-CoV-2 is a global pandemic that threatens to bring long-term changes worldwide. Approximately 80% of infected patients are asymptomatic or have mild symptoms such as fever or cough, while rest of the patients have varying degrees of severity of symptoms, with 3-4% mortality rate. Severe symptoms such as pneumonia and Acute Respiratory Distress Syndrome can be caused by tissue damage mostly due to aggravated and unresolved innate and adaptive immune response, often resulting from a cytokine storm. However, the mechanistic underpinnings of such responses remain elusive, with an incomplete understanding of how an intricate interplay among infected cells and cells of innate and adaptive immune system can lead to such diverse clinicopathological outcomes. Here, we use a dynamical systems approach to dissect the emergent nonlinear intra-host dynamics among virally infected cells, the immune response to it and the consequent immunopathology. By mechanistic analysis of cell-cell interactions, we have identified key parameters affecting the diverse clinical phenotypes associated with COVID-19. This minimalistic yet rigorous model can explain the various phenotypes observed across the clinical spectrum of COVID-19, various co-morbidity risk factors such as age and obesity, and the effect of antiviral drugs on different phenotypes. It also reveals how a fine-tuned balance of infected cell killing and resolution of inflammation can lead to infection clearance, while disruptions can drive different severe phenotypes. These results will help further the case of rational selection of drug combinations that can effectively balance viral clearance and minimize tissue damage simultaneously.

**Significance Statement:** The SARS-CoV-2 pandemic has already infected millions of people, and thousands of lives have been lost to it. The pandemic has already tested the limits of our public healthcare systems with a wide spectrum of clinicopathological symptoms and outcomes. The mechanistic underpinnings of the resultant immunopathology caused by the viral infection still remains to be elucidated. Here we propose a minimalistic but rigorous description of the interactions of the virus infected cells and the core components of the immune system that can potentially explain such diversity in the observed clinical outcomes. Our proposed framework could enable a platform to determine the efficacy of various treatment combinations and can contributes a conceptual understanding of dynamics of disease pathogenesis in SARS-CoV-2 infections.

## Introduction

The clinical phenotypes observed in the wake of the current COVID19 pandemic, caused by novel coronavirus SARS-CoV-2, are diverse. Nearly 80% of patients infected by the virus are either largely asymptomatic or show mild to moderate symptoms such as fever or cough. However, in a subset of patients, symptoms such as shortness of breath or pneumonia develop, thus requiring subsequent hospitalization. In a small percent of cases, severe pneumonia can aggravate to Acute Respiratory Distress Syndrome (ARDS), septic shock and multiple organ failure (1). Furthermore, there is diversity in the degree of ARDS and/or other pathological conditions observed within hospitalized patients; some, but not all, of them are admitted to Intensive Care Units (ICU) and require the support from mechanical ventilators. Higher age and/or pre-existing conditions such as diabetes or hypertension can serve as other comorbidities, thus influencing the severity of the disease and posing a higher risk of death to patients (2).

SARS-CoV-2 infection activates both innate and adaptive immune responses. These responses are responsible for viral clearance. In addition to neutralizing and eliminating the virus, the immune response may cause harmful tissue damage at both the local and systemic levels, when the regulatory mechanisms are dysfunctional. In a subset of individuals infected with SARS-CoV-2 dysregulated immune responses are observed, manifesting as a cytokine storm (3–5), which is reminiscent of immunopathology seen in other corona-viruses SARS-CoV and MERS-CoV (6). These observations suggest that the degree of dysregulation of immune response is likely to be an important factor, in addition to viral load, in determining the severity of COVID-19 (7, 8).

SARS-CoV-2, just like the SARS-CoV, attacks and enters host cells that harbour the ACE2 receptor. Serine protease TMPRSS2 is also required for viral entry for the S (spike) protein priming (9). ACE2 and/or TMPRSS2 have been seen to be present in multiple organs – lung, liver, colon, kidney, gut and placenta among others (10–13) making these organs potential targets of the SARS-CoV-2. Preliminary evidence suggests that similar to SARS-CoV (14), SARS-CoV-2 primarily targets type II alveolar cells (15, 16); thus, lung remains the primary organ of infection. Thus, the extent of lung injury has been proposed as a predictor of disease severity (17), possibly contributing to patient mortality with/without any co-morbidities (16).

With no specific cure identified for SARS-CoV-2, investigating the emergent dynamics of the interplay between the infected cells and the components of innate and/or adaptive immune system is likely to help understand the different immunopathology scenarios, and devise effective therapeutic strategies that can curb the disease severity. Mathematical models have been extensively used to describe the dynamics of viral load in patients for HIV, HCV, influenza and Ebola virus (18); however, the integration of immune system dynamics in those frameworks are relatively recent (19–21). In the context of SARS-CoV-2, a quantitative and predictive framework to understand the dynamics of different possible outcomes of the complex nonlinear interactions among infected cells and immune system remains to be constructed.

Here, we have developed a mathematical model to capture the interconnected dynamics of infected cells and the components of innate and adaptive immune system. Our model recapitulates the spectrum of outcomes observed in SARS-CoV-2 patients – from being asymptomatic to different severe manifestations of the disease via increased cytokine-mediated immunopathology. Thus, we offer a framework that can be used to both explain the contribution of different host-virus interactions to augmenting disease severity, as well as to serve as a predictive platform to identify mechanisms to limit viral invasion of the host tissue as well as the consequent immunopathology, hence contributing to the existing knowledge base to reduce the disease burden of the ongoing pandemic.

## Results

### Innate immunity system can lead to diverse outcomes for viral clearance and immunopathology

The host innate immune response is the first line of defense once the virus enters the body (22). Thus, as a first step, we model the following set of interactions among the infected cells (V) and the innate immune system (M) (**Fig 1A**). First, SARS-CoV-2 virus can replicate in infected cells and consequently infect other healthy cells in a logistic growth fashion (rate: *kv*). Second, the virus itself or the infected cells can elicit an innate immune response via pattern recognition receptors expressed on the immune cells or through the production of cytokines and chemokines that can attract other cells of innate immune system such as neutrophils, natural killer cells, monocytes and macrophages (rate: *kvm*) (23). For the purpose of simplicity, we have combined the response of all these individual innate immune cell populations into one entity represented as innate immune cells (M). Third, innate immune system cells can induce recruitment and/or proliferation of other cells of innate immune system via the production of cytokines or chemokines (rate: *kmm*), enabling a self-sustaining response in the presence of infected cells (24). Fourth, infected cells and thereby the virus can be cleared by innate immune cells (rate: *dmv*) (25, 26). Fifth, in the absence or at very low levels of infected cells, either due to the clearance of the virus or under homeostatic conditions, accumulation of innate immune cells can be self-suppressing (rate: *dvm*) (27, 28).

**Fig 1:**
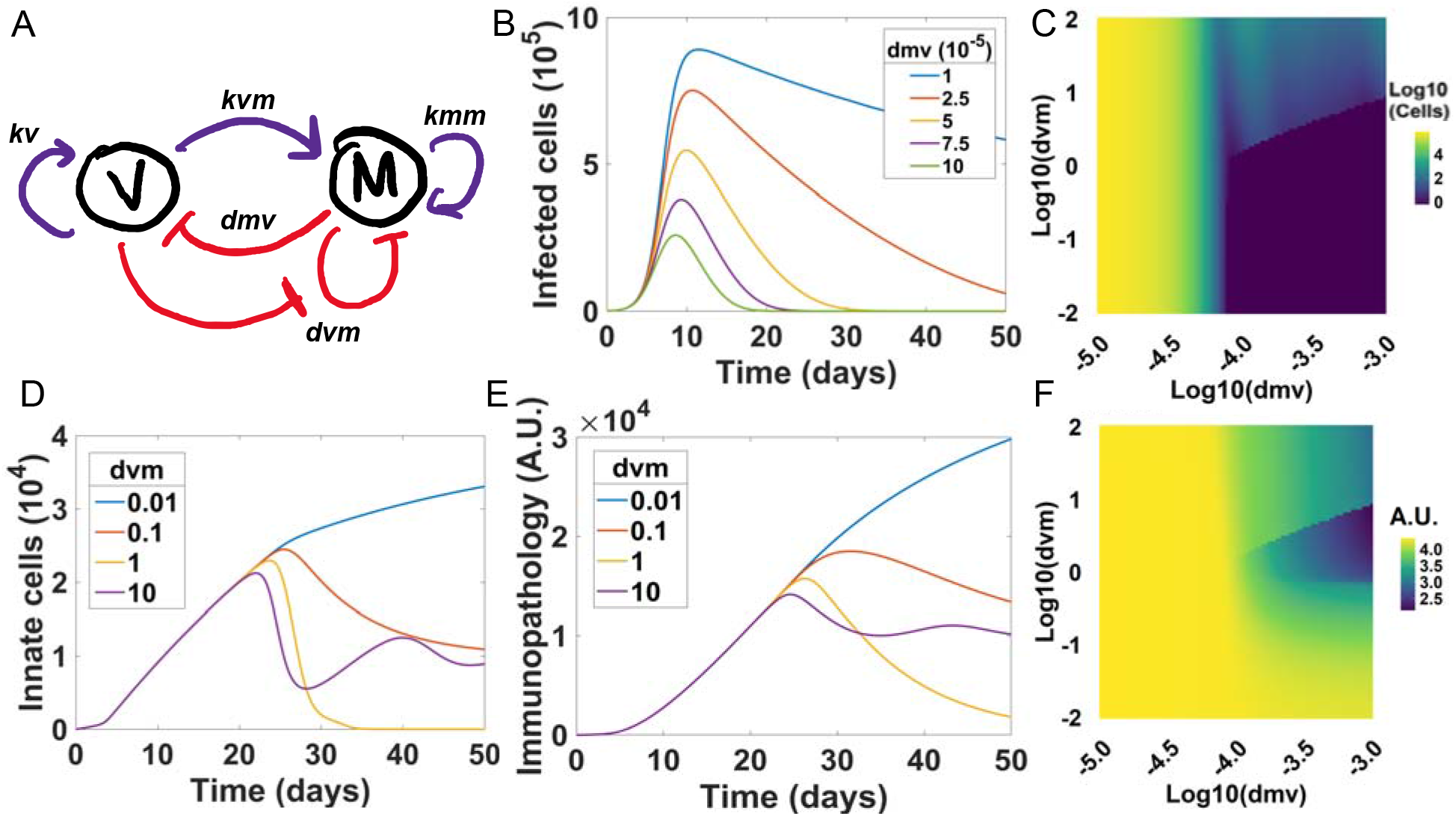
Dynamics of interaction among infected cells and innate immune system cells. **A)** The network representing the interactions between infected cells and innate immune cells. **B)** Dynamics of infected cell numbers for varying values of dmv (10e-5 to 10e-4). **C)** Effect of dvm and dmv on infected cell load at 30 day time point. **D)** Effect of dvm on the dynamics of innate immune cells. **E)** Effect of dvm on immunopathology. **F)** Effect of dvm and dmv on immunopathology at 30 days.

To elucidate the dynamics of these interactions between the infected cells and the innate immune system, we modeled this system using a system of ODEs (ordinary differential equations) **(Supplementary Methods)**. We observed that the clearance time of the virus decreases drastically with an increased killing rate of infected cells via the innate immune cells (*dmv*) (**Fig 1B**), suggesting that the initial response mounted by the body to the virus might be paramount in resolving the infection at an early stage. This trend is qualitatively maintained for other values of *kv* (**Fig S1A**). The parameter *kv* can be considered as a representation for net infectivity of the virus, which can be a result of the replication rate of the virus inside the host (for instance, *kv* can be decreased by intra-cellular interferon response in infected cells (29)), its exit efficiency from the cells, its survival outside host cells and the viral infection rate of neighboring healthy host cells. Therefore, as expected, the higher the rate of growth of infected cells *kv*, the higher the peak of infected cells and the more is the time required to clear the infection (**Fig S1B**). Thus, these results suggest that some strains of the virus that may have an inherent lower infection/proliferation/survival rates can get cleared more efficiently than those that are highly virulent.

Sensitivity analysis of the system for the time of clearance of virus (time till V < 1) reveals that the growth rate of infected cells (*kv*) and the killing rate of infected cells via the innate immune cells (*dmv*) are crucial in controlling the infection within a stipulated time (**Table S1**), further underscoring the relevance of these parameters in controlling the infection at an early stage. Viral clearance times were found to be positively correlated with *kv* and inversely correlated with *dmv* (**Fig S1C**).

Immunopathology is a crucial parameter that defines the severity of the disease that the patients suffering from SARS-CoV-2. To that extent, we defined immunopathology (P) by the following equation, primarily with the goal of capturing its cytokine-mediated effects:

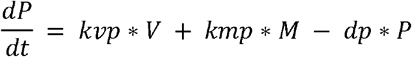

where P is the immunopathology in the system, *kvp* is the rate at which infected cells (V) contribute to the immunopathology by secreting pro-inflammatory cytokines such as IL-6 *etc.* (23), *kmp* is the rate at which innate immune cells (M) contribute to immunopathology due to creation of a strong pro-inflammatory environment (cytokine storm) (30), and *dp* is the rate at which the system recovers from the immunopathology following a first order decay (19).

We hypothesized that the rate of resolution of innate immune cells via self-suppression as the number of infected cells decrease (*dvm*) is a crucial parameter that affects immuno-pathology in the system. To test this hypothesis, we monitored the infected cell population in the system at the end of 30 days as a function of both *dmv* and *dvm* (**Fig 1C**). It is expected that the higher the *dmv*, the smaller the number of infected cells at the end of 30 days, irrespective of the value of *dvm*. One would expect that the rate of resolution of innate immune system (*dvm*) will minimally affect the levels of infected cells, because the resolution is thought to be triggered after the clearance of infected cells and hence has very little role to play in the clearance itself. This fact holds true for low values of *dvm*, but surprisingly not for higher values of *dvm* (**Fig 1C**). At high values of *dvm*, some viral load (in the form of infected cells) persists, while there is complete clearance at lower values of *dvm* (**Fig S1D**). It is, however, important to note here that the peak number of virally infected cells depends primarily on *kv* and *dmv* (**Fig S1E**) but not on *dvm* (**Fig S1F**). This dynamics may be one among many contributing factors to persistence of the virus as seen in other diseases such as HCV (31). The persistence of virally infected cells can lead to and/or is a result of the concurrent maintenance in the levels of innate immune cells (**Fig 1D**) which can consequently give rise to varying levels of immunopathology (**Fig 1E**). Finally, we tracked the immunopathology levels at the end of 30 days and observed that although the parameter *dvm* does not affect the immunopathology at lower values *dmv*, it has a non-monotonic relation at higher *dmv* values with intermediate values of *dvm* giving the least immunopathologic conditions **(Fig 1F)**. It is also important to note that although the final immunopathology at the end of 30 days shows a dip at intermediate values of *dvm*, such non-monotonic responses are not seen in the peak immunopathology value either in a two-dimensional scan of *kv*-*dmv* or *dmv*-*dvm* (**Fig S1G-H**). Put together, these observations suggest that initial innate immune response can drive varying outcomes in terms of viral clearance and immunopathology, with a possibility of viral persistence.

### Adaptive immune system can boost the clearance of infected cells resulting in ‘non-severe’ phenotypes

Next, we introduced adaptive immune system dynamics into the system as well (**Fig 2A**) by considering following additional interactions. First, cells of the adaptive immune system, primarily the cytotoxic CD8^+^ cells, tend to proliferate in an antigen dependent manner in proportion to viral load (denoted here by the number of infected cells) (19). We model this growth of adaptive immune cells in a logistic fashion but also dependent on the number of infected cells. Second, these adaptive immune cells can kill the infected cells at a rate of *dcv*. Third, the infected cells can induce exhaustion in the adaptive cell population, reducing the number of functionally active adaptive cells to kill the infected cells, through constant antigen accumulation, when present in overwhelmingly large numbers (32, 33). Furthermore, adaptive immune cells (C) are also known to contribute to immunopathology (34, 35), so we add a new contribution term *kcp*\**C* to the previous equation where *kcp* is the rate at which adaptive cells contribute to the immunopathology. The consequent equation thus becomes:

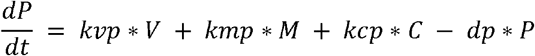

**Fig 2:**
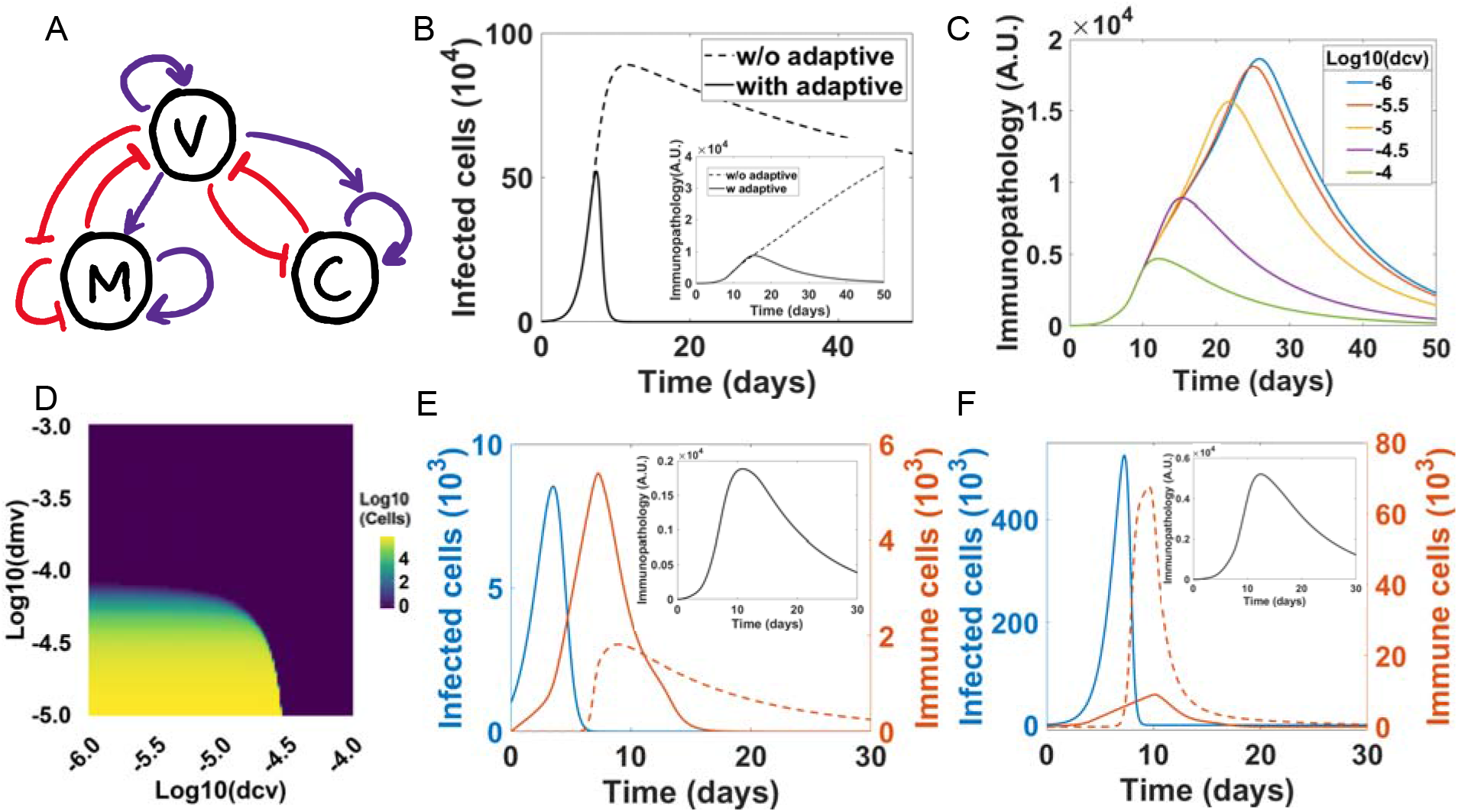
Adaptive immune system aids the innate immune system. **A)** Network representing the interactions between infected cells, innate immune cells, and adaptive immune cells. **B)** Demonstration of improved clearance of infection upon induction of an adaptive immune response (solid line) as compared to the case without it (dotted line). Inset shows the reduction of immunopathology upon induction of adaptive immunity. dmv= 1e-5, dcv:0 (w/o adaptive), 5e-5 (with adaptive). **C)** Effect of dcv on immunopathology. **D)** Heatmap showing the infected cell count 30 days post symptom onset for varying values of dcv and dmv. **E)** Infected and immune cell profiles in case of a strong innate immune system (healthy phenotype 1). dmv=1e-3, dcv=1e-6. Inset shows the immunopathology values. **F)** Same as E) but for a strong adaptive immune system (healthy phenotype 2). dmv=1e-6, dcv=1e-4. For the figures E and F, the solid orange line represents innate immune cell population while the dashed line represents adaptive immune cell population.

In cases where the innate immune system was unable to clear the virus (for example, low dmv denoted by blue curve in **Fig 1A**), introduction of the adaptive immunity achieves viral clearance (**Fig 2B**) and can help reduce corresponding immunopathology (**Fig 2B inset**). With the effective clearance of the virus by the adaptive immune cells, no buildup of either the innate or adaptive immune system population is seen (**Fig S2A**). Furthermore, the strength of clearance of the virally infected cells by adaptive immune cells (*dcv*) influences the levels of immunopathology. The higher the value of *dcv*, the faster the clearance rate (**Fig S2B**) and the smaller the immunopathology peak value (**Fig 2C**).

We then investigated how the number of infected cells depend on the relative strengths of the components of innate and adaptive immune system - *dmv* and *dcv* - correspondingly (**Fig 2D**). High degree of effectiveness by either of the immune system arms was sufficient to clear the number of infected cells in a period of 30 days, thus offering a possible explanation for a majority of SARS-CoV-2 patients being asymptomatic. However, if the efficacy of both the arms of the immune system is low, the virus is not cleared efficiently, and consequent immunopathology values are higher (**Fig S2C**). Furthermore, the peak of infected cells depends more on the values of *dmv*, pointing to the fact that the strength of the innate immune system is perhaps more crucial in determining the peak infected cells compared to adaptive immune system (**Fig S2D**).

Although either of the two arms of the immune system may be sufficient to clear the infected cells, it must be noted that these two arms may be contributing to varying degrees in cases of patients falling in this broad clinical phenotype of “non-severe” (**Fig 2E-F, S2E**). Thus, perturbing these dynamics can have a patient-specific response. We specifically considered two possible extreme cases: a very strong innate immune system but a very weak adaptive immune system and *vice versa*. In the “non-severe” case shown in **Fig 2E**, death rate of infected cells due to killing by the innate immune system (*dmv*) is 3 orders of magnitudes higher than that of the adaptive immune system (*dcv*), and vice versa in the case shown in **Fig 2F**. These differences cause some marked differences in the modes of clearance of infected cells from the system. As observed, the peak number of virally infected cells is far lower when the innate arm is stronger (**Fig 2E**). This observation can be partly explained by the fact that the innate immune system is acting much earlier than the adaptive immune system. Interestingly, the viral clearance in both cases happens within 15 days, concordant with clinically observed data of viral clearance within 14 days in completely asymptomatic cases (36).

Interestingly, the peak levels of immunopathology are slightly (i.e. three-fold) higher in the case when the adaptive immune system is relatively more effective than the innate immune (compare insets in **Fig2F** and **Fig 2E**). This difference can be partly explained by a slightly larger accumulation of innate immune cells, which contribute to immunopathology more than the adaptive immune cells, because the adaptive system kicks in much later as compared to innate immune system. This remnant immunopathology scenario (**Fig 2F**) may manifest as mild symptoms within the broad definition of “non-severe” cases, for instance in the form of fever, cough, and/or breathlessness, whereas the other scenario (**Fig 2E**) may be completely asymptomatic. As expected, the exhaustion of adaptive cell activity is higher in case of stronger adaptive response **(Fig S2F).** While we have presented two extreme scenarios here as representative examples, in reality, there is likely a continuum of such phenotypes with varying degrees of contributions from the two arms in immune system that can give such “non-severe” phenotypes.

### Clinically observed ‘severe’ phenotypes of COVID-19

To test the hypothesis that the more “severe” SARS-CoV-2 mediated phenotypes can possibly arise in cases of inefficient resolution of the inflammation caused by the innate immune system (**Fig 1D-E**), we reduced the value of *dvm* by ten-fold. Such a reduction would imply the set of cases where the resolution of inflammation by the innate immune system is inefficient. Such sustained innate immune system has been seen to be specific to non-survivors in SARS-CoV, suggesting that “being stuck in immune response” can lead to a severe disease phenotype (37). This lack of ability to resolve innate immune response and mount an efficient adaptive response is often an age-related change; this reason may at least in part explain higher susceptibility to the severe phenotype for elder population, ultimately leading to their death (37). Similarly, people with comorbidities such as obesity have heightened inflammatory levels at baseline (in the absence of infection), which could affect resolution in cases of inflammation (38). For this scenario with reduced *dvm*, we observed that similar to the previous scenario with a higher value of *dvm*, effective clearance of infected cells was possible as long as a strong innate and/or adaptive immune system arm is functional (**Fig S3A**). However, the extent of immunopathology is drastically higher in this case than in the previous case (compare **Fig S3B** with **Fig 2C**). Clearly, the lower resolution strength of the innate immune system is pushing the system from a lower immunopathology regime to a higher one. However, it is important to note here for most parametric combinations of *dmv* and *dcv*, infected cells are still ultimately cleared form the system. This observation is in concordance with observations that the deaths/complications in this pandemic is not likely to be primarily due to the inability of the system to clear the virus, but due to the unresolved immune responses.

Interestingly, we found that in this parametric regime of a lower value of *dvm*, many “severe” phenotypes can be observed. First, when the adaptive immune system is relatively strong/robust, the infection is cleared efficiently from the system (**Fig 3A**) at a similar timescale (<10-15 days) as noticed in previous “mild” scenario (**Fig 2F**). However, a sustained level of the innate immune cells (mostly macrophages) are retained, leading to immunopathology (**Fig 3B**). The level of immunopathology is higher than that observed for the mild scenario (**Fig 2F inset**), but lower than many other severe phenotypes mentioned later. Thus, this phenotype may map on to the subset of patients who suffer from “pneumonia-like symptoms” and hence require hospitalization but most likely will not require admission to intensive care unit (ICU).

**Fig 3:**
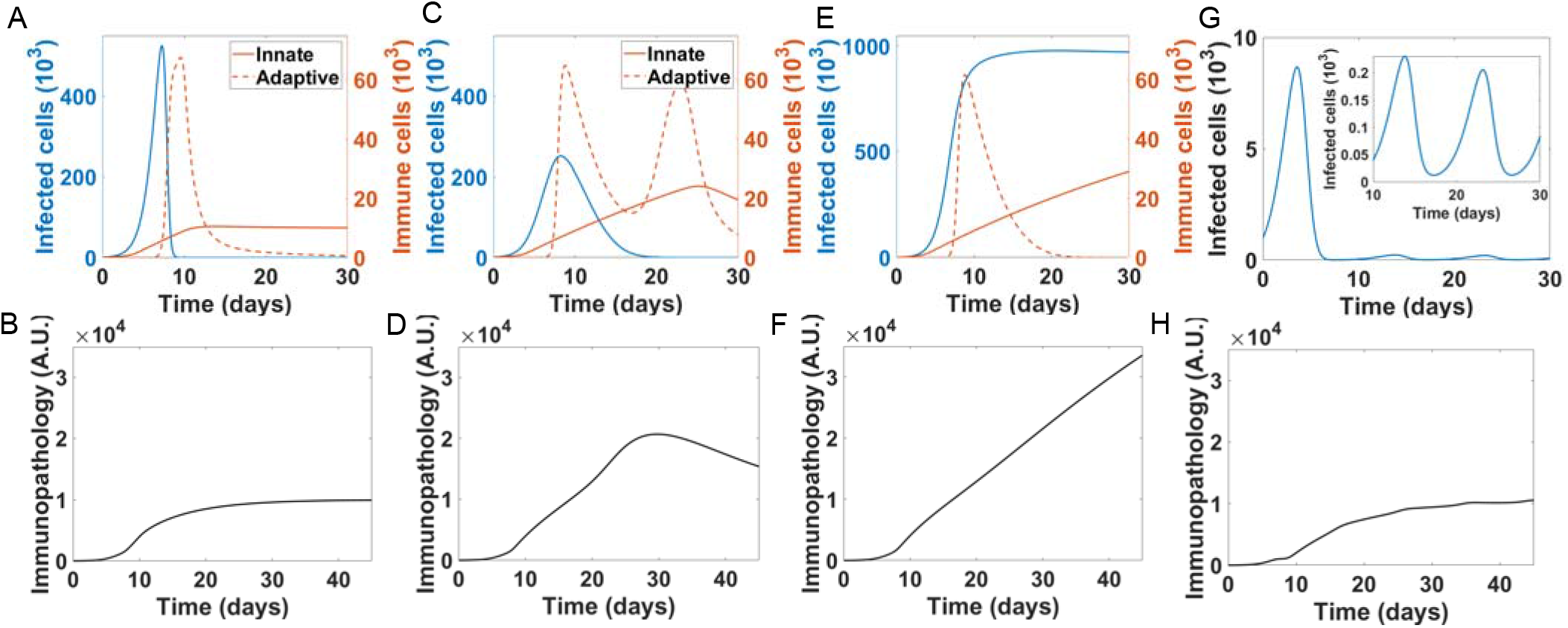
Clinically observed phenotypes of COVID-19. **A)** Diseased counterpart of the healthy phenotype in Fig. 2F; reduced dvm leads to persistent innate immune cells (severe phenotype 1). dmv= 1e-6, dcv = 1e-4, dvm = 0.1. **B)** Dynamics of immunopathology for case shown in A. **C)** Similar to A but corresponding to a case represented by Fig. 2E (severe phenotype 2). dmv = 1e-4, dcv = 1e-6, dvm = 0.1. **D)** Dynamics of immunopathology for case shown in B. **E)** Persistent immunopathology accompanied by lack of infection clearance due to a weak innate and adaptive immune system (severe phenotype 3). dmv =1e-6, dcv = 1e-6, dvm =0.1. **F)** Dynamics of immunopathology for case shown in E. **G)** Infection persistence observed for high dvm. Inset shows the sustained infection via oscillations. dcv= 1e-6, dmv= 1e-4, dvm= 10. **H)** Immunopathology corresponding to the case shown in G.

In contrast to the above phenotype, we observe a slightly different phenotype **(Fig 3C)** where the infected cell clearance takes a relatively longer time (15-20 days). This phenotype occurs with approximately two-fold higher levels of immunopathologic damages than that the previous case (**Fig 3D** vs **Fig 3B**). Such patients might require admission into the ICU, but their outcome depends on the degree of immunopathology the body can handle. In excellent match with our model predictions, recent clinical data suggests that the COVID-19 patients admitted to the ICU have a longer viral clearance time than the non-ICU patients: nearly 30 days vs. 12-16 days (39). Furthermore, the adaptive cell population is far more exhausted in this scenario as compared to the previous “pneumonia-like symptoms” case (**Fig S3E:** blue curve vs orange curve), again reminiscent of clinical observations of severe cases of COVID-19 patients (40).

Another phenotype in which the infection spread in the body becomes uncontrollable is when the efficacy of both innate and the adaptive immune arms is low. In such scenario, sustained high levels of the virus and infected cells in the body without any clearance at all **(Fig 3E)**. We posit that this scenario is the most severe of all phenotypes; the immunopathology levels are high and constantly increase with time **(Fig 3F vs Fig 3B or 3D).** Under such circumstances, the levels of the exhaustion rates seen in the adaptive immune cells will be the most (**Fig S3E**: yellow curve). This scenario is most likely to be fatal for the patients with a high viral load at the time of death (41), unless there are aggressive intervention strategies to lower the virus and reduce strong pro-inflammatory environment as well.

After investigating these three different severe phenotypes which were obtained at a ten-fold lower *dvm*, we probed the behavior of the system at a ten-fold higher *dvm* value. Higher *dmv* value may lead to persistence of infection in the system, because the self-suppression of innate immune system kicks in quite strongly which can further impede the clearance, unless higher values of *dcv* are brought in (compare **Fig S3F** with **Fig S3A**). Thus, in the presence of strong self-suppression of innate immune system, a strong adaptive immune system seems to be necessary and sufficient to clear the infection. Consequently, in the subset of patients who have a relatively strong resolution capacity of the innate immune system, but a weaker adaptive system will tend to harbor a persistent population of the infection with moderate levels of immunopathology (**Fig S3G**). Particularly for the scenario of relatively high values of *dmv* as compared to *dcv* for which the infection clearance happens in 15-20 days (**Fig 3C**), a ten-fold increase in *dvm* renders the levels of virally infected cells to be oscillatory, albeit at lower load (approximately, 100 infected cells in the system) (**Fig 3G**). This phenotype gives intermediate values of the immunopathology (**Fig 3H**), with innate and adaptive immune cell count kept in check (**Fig S3H**). This phenotype may correspond to a subset of patients who are likely to have a prolonged stay at hospital (>30 days) but without overwhelmingly “severe” symptoms. Put together, these observations point towards the fact that optimal immune resolution responses are crucial in determining the “severe” clinical phenotype outcome in SARS-CoV-2 infections.

### Effect of virulence reducing interventions on the clinical phenotypes of COVID-19

Most antiviral drugs such as remdesivir aim to reduce the virulence of the infection by inhibiting in-host viral reproduction, thus enhancing the chances of clearance via the immune system (42). Because the severity of clinical phenotypes of COVID-19 is majorly due to immunopathology and our model suggests a complex non-linear relationship between immunopathology and viral clearance, we investigated how the reduction of virulence effects the clinical phenotypes in COVID-19. Specifically, we analysed the effect with respect to two intervention parameters, intervention time (T) (number of days after viral infections when the drug is first administered) and the efficacy of the intervention, defined by the percentage reduction in the parameter *kv* upon drug administration. For simplicity, we assume that *kv* remains reduced to a constant value after the administration of the drug.

For the parameter set that had a moderately strong innate immune system but impaired adaptive immune system and enabled viral clearance in about 20 days (Phenotype shown in **Fig 3C**), intervention at 10 days with 50% efficacy reduced the infection clearance time and peak immunopathology achieved (**Fig 4A**). In addition, under these conditions, the peak load of innate immune cells also reduced while having minimal changes in the peak values of adaptive immune cells (**Fig 4B**). We next quantified the effect of drug treatment on these parameters by altering the time at which the antiviral therapy was introduced. We observed that there is a considerable reduction in the immunopathology peaks observed, with the maximum difference being caused for intervention time around 10 days, right after the infection peak (**Fig 4C**); this difference scaled with the efficacy of treatment introduced. However, there was significant lowering in the infection clearance times (**Fig 4D**) and the peak infected cell values, if the intervention was made earlier than 10 days (**Fig S4A**). These results emphasises the fact that even though one were to reduce the viral load in the system by giving antivirals, it might not be sufficient to bring down the immunopathology related complications without the use of an additional drugs that alter immune activity.

**Fig 4.**
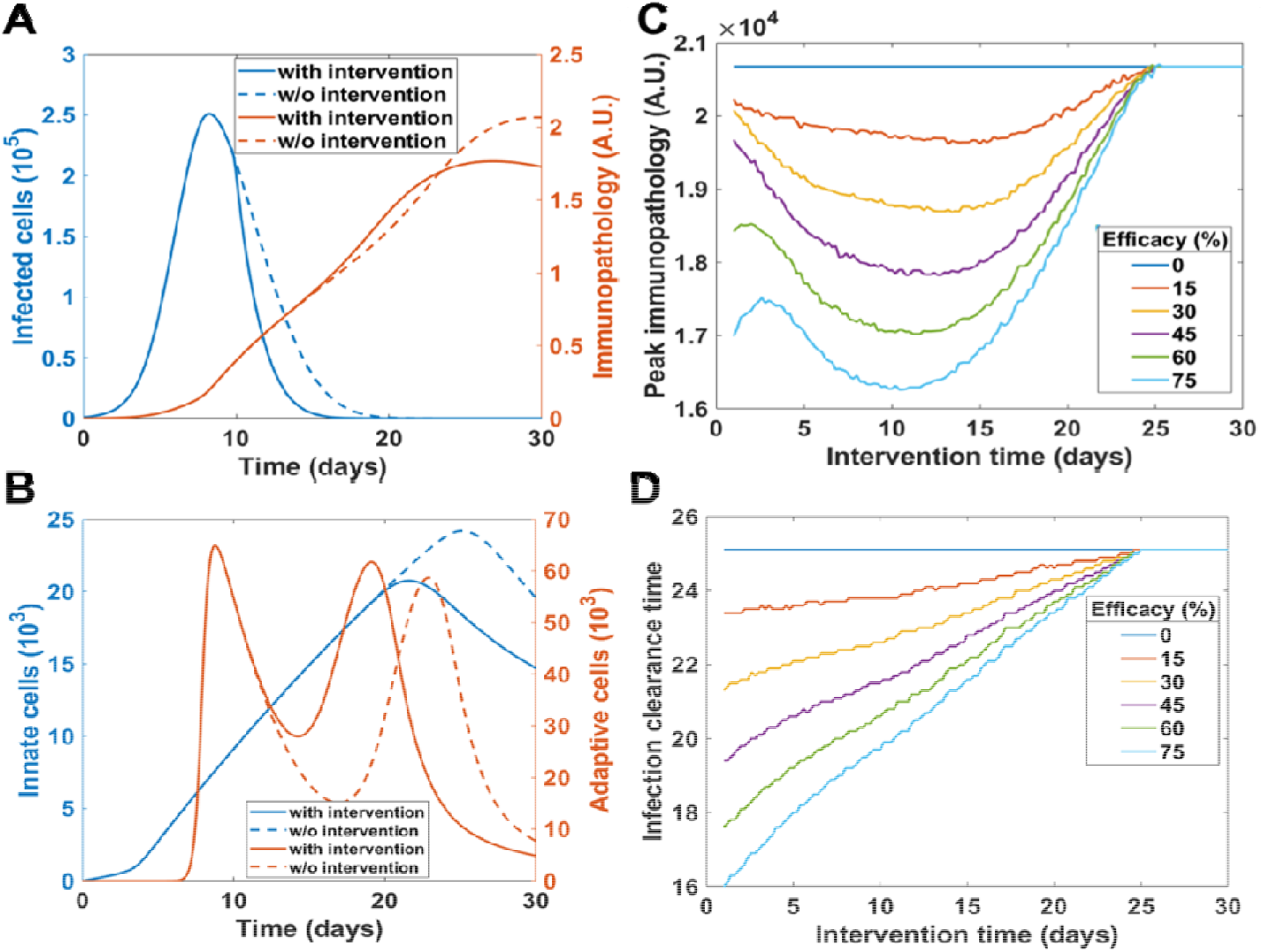
Effect of antiviral intervention on viral clearance and immunopathology. A) Dynamics of infected cell and immunopathology with (50% efficacy) and without (0% efficacy) drug intervention at 10 days. **B)** Immune cell profiles for the case in A. **C)** Effect of intervention time and drug efficacy on peak immunopathology for the phenotype in Fig 3C-D. Parameters taken from Fig 3C-D. **D)** Same as A, but for infected cell peak

Furthermore, we performed the same analysis for the higher immunopathology phenotype seen earlier (**Fig 3E**) and observed that the efficacy of antiviral treatment in reducing the peak viral load or decreasing the infection clearance time was minimal (**Fig S4B,S4F**). However, the effect of antiviral treatment was more pronounced in the scenario of lower immunopathology scenario (**Fig 3A**) in terms of both peak viral load (**FigS4D**) and enabling faster clearance (**Fig S4B**). Indeed, the peak of infected cell count does decrease with increasing drug efficacy, as *kv* directly affects the rise of infection levels. We see these effects diminishing as the intervention time reaches closer to the time taken for infected cells to reach the peak. These efforts, however, had little to no effect in reducing the immunopathology levels significantly (**Fig S4C, S4E**). These results suggest that the antiviral drugs targeting the virulence of infection may not be equally effective in reducing immunopathology in all the “severe” phenotypes in COVID-19 patients. The impact of these drugs is likely to be maximal for the patients with a relatively weak adaptive and relatively strong innate immune system. This aspect also underscores the importance of using immune suppressants like IL-6 receptor antagonists to simultaneously decrease viral load and reduce immunopathology and facilitate tissue repair.

### Effect of regulatory T cells on clinical phenotypes of COVID-19

Regulatory T cells have been shown to be essential modulators of innate immune system in maintaining tissue homeostasis especially during various respiratory disorders (43). They can limit immune inflammation-inflicted tissue damage and facilitating tissue repair, accelerating the resolution of acute lung injury (ALI) (44). However, increased presence (or activity) of regulatory T cells may end up suppressing the immune responses that are important for viral clearance, and consequently cause viral persistence (45). Thus, we examined the role that regulatory T cells play in the context of SARS-CoV-2.

In our model, we included a node representing such regulatory T cells that can be activated by the virally infected cells in an antigen-dependent manner (**Fig 5A**). These regulatory cells can directly inhibit the function of innate (43) and adaptive (46) immune system largely via secreting IL-10 or TGF-β. We observed that parameter sets with robust viral clearance but high immunopathologic levels were now able to resolve the immunopathology with the introduction of the regulatory T cells without necessarily a change in the dynamics of virally infected cells **(Fig 5B)** but with a subsequent reduction in the innate immune cell levels in comparison to the control case (compare **Fig S5A, S5B**). An extremely strong regulatory cell response can lead to over-suppression of the innate and the already impaired adaptive immune system (compare **Fig S5C, S5D**) which may result in the resurgence of the infection (**Fig 5C)**. Furthermore, a strong immune suppression of both the arms of the immune responses by the regulatory cells causes an apparent clearance of the virus but the virus can persist at very low levels until the immune cells are suppressed enough (compare **Fig S5E, Fig S5F**) for the resurgence of the virus **(Fig 5D).** This scenario of immuno-compromised state can allow other opportunistic pathogens to infect the lungs (47), thus further complicating the situation. These analyses suggest that although the regulatory T cells have the capability of resolving cytokine mediated immunopathology, overactivation of the same can enhance non-clearance of the virus and facilitate other infections.

**Fig 5.**
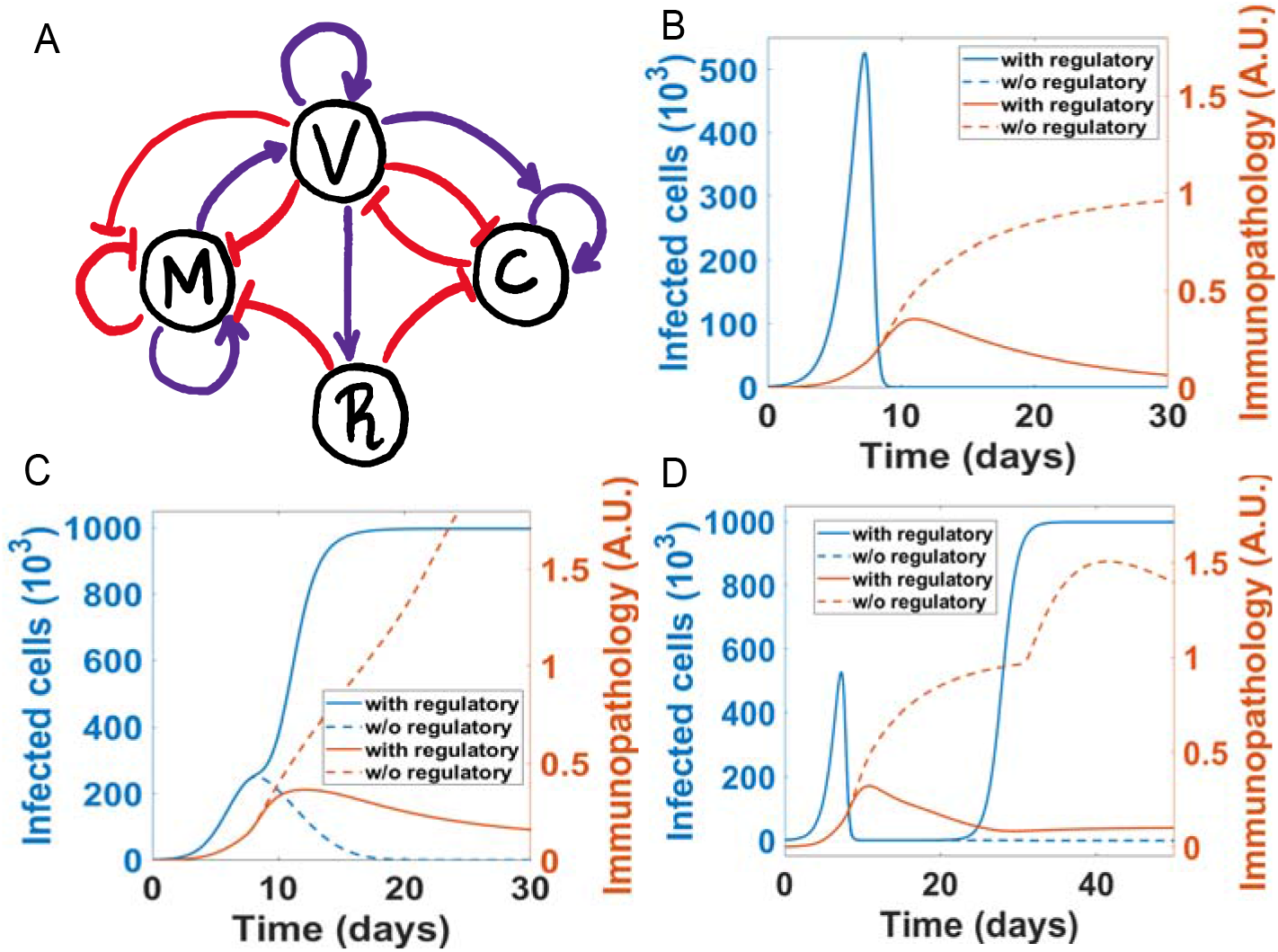
Effect of regulatory cells on clinical phenotypes. **A)** Network representing the interactions between infected cells, innate and adaptive immune cells and regulatory cells. **B)** Reduction of immunopathology by regulatory cells in severe phenotype 2 (Fig 3C-D) with no effect on infection profiles. **C)** Reduction of immunopathology accompanied by non-clearance of infection for severe phenotype 1 (Fig 3A-B). **D)** Strong regulation of immune cells by regulatory cells leads to persistence and resurgence of infection. Parameters for B: dmv = 1e-6, dcv = 1e-4, dvm = 0.1, drm = 0 and 1e-4, drc = 0 and 1e-5. Parameters for C: dmv = 1e-4, dcv = 1e-6, dvm = 0.1, drm = 0 and 1e-4, drc = 0 and 1e-5. Parameters for D: dmv =1e-4, dcv = 1e-6, dvm = 0.1, drm = 0 and 1e-4, drc =0 and 1e-4.

## Discussion

Immune reactions, which are initiated in response to a SARS-CoV-2 infection but turn aberrant, are increasingly being implicated in the complications and deaths associated with SARS-CoV-2 (48–50). However mechanistic underpinnings of such dysregulation remain largely underexplored. Mathematical modeling efforts for COVID-19, similar to the case for other infectious diseases (51), have mainly focused on epidemiological dynamics (52, 53), relative to those focusing on within-host dynamics (54, 55). Such intra-host dynamics models have been extensively studied for HIV, HCV and cancer (56–59).

Here, we attempt to offer mechanistic insights into existing empirical observations on diverse phenotypes in SARS-CoV-2 infections with the already established mechanistic links between the components of the immune system and their interactions of virally infected cells. We have developed a minimalistic yet rigorous mathematical model that encapsulates the interactions between the virally infected cells and the innate and adaptive immune systems to observe the various clinically observed phenotypes emerge as a result of the dynamics of the system. This mathematical model uncovers the existence of distinct phenotypes and parametric combinations that enable them. Thus, we uncover the delicate balance that exists between viral clearance, immunopathology and resolution of inflammation and how the various combinations of these give rise to some of the observed phenotypes.

Our model, within biologically reasonable parameter ranges, closely mimics some of the observed clinical patterns. First, our model closely follows the observed trends of viral loads peaking around 5-6 days after symptom onset (60). In our model, virally infected cells which act as a proxy for viral load in the respiratory system of a patient peak around 2-10 days. Second, the peak of infected cells in severe vs. non-severe (mild/asymptomatic) phenotypes in our model does not differ largely, reminiscent of clinical observations that the observed viral load is similar for symptomatic vs. asymptomatic patients (8). Third, our model predicts that patients with higher immunopathology values are associated with longer infection clearance times, a prediction that is validated by recent clinical data suggesting that COVID-19 patients admitted to the ICU tend to take twice the clearance time compared to the non-ICU patients (30 days vs. 14 days) (39). Fourth, we demonstrate that the impaired ability of innate immune system to resolve the inflammation can drive cytokine-induced immunopathology; indeed, “being stuck in immune response” can be a hallmark of severe disease and possibly the death of the patient (37). Similar observations have been made for SARS-CoV and MERS-CoV (61); for instance, patients who died of SARS had higher levels of pulmonary pro-inflammatory cytokines (62). Thus, while our model is relatively coarse-grained and clubs many innate and adaptive innate immune systems into one variable each, it is capable of recapitulating various clinicopathological features for COVID-19 patients reported so far.

Future work will involve a more detailed agent-based modeling integrating the intracellular signaling mechanisms with the cytokine-mediated paracrine crosstalk among different cell types that may get infected and the components of innate and adaptive immune system that operate in different tissues. While the viral entry mechanisms may be similar for SARS-CoV and SARS-CoV-2, the similarities and differences in impact of internalized virus on intracellular signaling in infected cells seems to be yet identified. Interestingly, SARS-CoV was shown to be able to suppress the interferon response through viral proteins, hence overcoming one of the first lines of defense of the body (63–65). Preliminary evidence suggests that the SARS-CoV-2 can induce a robust expression of interferon pathway related genes (12, 66). However, the degree of interferon response elicited by individual patients can vary. Our model parameter k_v_ can capture variability in this feature, and can be incorporated as a function of underlying signaling pathways that drive this variability, such as NF-kB signaling (67, 68).

Currently, it is still unclear for SARS-COV-2 how the viral proteins might interfere in the normal functioning of the immune system of the body. Such crosstalk, once discovered, can be incorporated in the model. Given the diverse clinical phenotypes observed, we will likely need to design different therapeutic regimens. A recent study showed that in a cohort of 100 patients with hyperinflammatory syndrome, Tocilizumab, a monoclonal antibody that targets the IL6 receptor, can improve or stabilize the condition in about 77% of the cases, whereas the condition worsened in remaining 23% (69). This difference also points to the fact that in cases of hyperinflammatory cases which lack inflammation resolution strength, it might not be enough to give anti-virals only; instead, immune suppressants become a must as a combination therapy. However, such a therapy is likely to fail in phenotypes where there is a strong resolution of inflammation, leading to viral persistence. Thus, our model can serve as a decision-making and predictive framework in such situations where one needs to distinguish between different possible phenotypes and develop mechanistic drug combinations and optimize intervention time of these combinations.

## Author contributions

MKJ designed the research, MKJ and SJ supervised research, SS and KH performed research, all authors contributed to data analysis and writing of the manuscript.

## Acknowledgements

MKJ was supported by Ramanujan Fellowship provided by Science and Engineering Research Board (SERB), Department of Science and Technology, Government of India (SB/S2/RJN-049/2018).

## References

1. Richardson S, et al. (2020) Presenting Characteristics, Comorbidities, and Outcomes Among 5700 Patients Hospitalized With COVID-19 in the New York City Area. JAMA:in press.

2. Yang J, et al. (2020) Prevalence of comorbidities and its effects in coronavirus disease 2019 patients: A systematic review and meta-analysis. Int J Infect Dis 94:91–95.

3. Huang C, et al. (2020) Clinical features of patients infected with 2019 novel coronavirus in Wuhan, China. Lancet 395:497–506.

4. Ye Q, Wang B, Mao J (2020) The pathogenesis and treatment of the “Cytokine Storm” in COVID-19. J Infect 80:607–613.

5. Jose RJ, Manuel A (2020) COVID-19 cytokine storm: the interplay between inflammation and coagulation. Lancet Respir Med. doi:10.1016/S2213-2600(20)30216-2.

6. Channappanavar R, Perlman S (2017) Pathogenic human coronavirus infections: causes and consequences of cytokine storm and immunopathology. Semin Immunopathol 39(5):529–539.

7. To KKW, et al. (2020) Temporal profiles of viral load in posterior oropharyngeal saliva samples and serum antibody responses during infection by SARS-CoV-2: an observational cohort study. Lancet Infect Dis 20:565–74.

8. Zou L, et al. (2020) SARS-CoV-2 viral load in upper respiratory specimens of infected patients. N Engl J Med 382:1177–1179.

9. Hoffmann M, et al. (2020) SARS-CoV-2 Cell Entry Depends on ACE2 and TMPRSS2 and Is Blocked by a Clinically Proven Protease Inhibitor. Cell 181(2):271–280.e8.

10. Qi F, Qian S, Zhang S, Zhang Z (2020) Single cell RNA sequencing of 13 human tissues identify cell types and receptors of human coronaviruses. Biochem Biophys Res Commun 526(1):135–140.

11. Ashray N, et al. (2020) Single-Cell RNA-seq Identifies Cell Subsets in Human Placenta That Highly Expresses Factors to Drive Pathogenesis of SARS-CoV-2. Preprints:2020050195.

12. Lamers MM, et al. (2020) SARS-CoV-2 Productively Infects Human Gut Enterocytes. Science:eabc1669.

13. Hamming I, et al. (2004) Tissue distribution of ACE2 protein, the functional receptor for SARS coronavirus. A first step in understanding SARS pathogenesis. J Pathol 203(2):631–7.

14. Mossel EC, et al. (2008) SARS-CoV replicates in primary human alveolar type II cell cultures but not in type I-like cells. Virology 372(1):127–135.

15. Chu H, et al. (2020) Comparative replication and immune activation profiles of SARS-CoV-2 and SARS-CoV in human lungs: an ex vivo study with implications for the pathogenesis of COVID-19. Clin Infect Dis:ciaa410.

16. Li H, et al. (2020) SARS-CoV-2 and viral sepsis: observations and hypotheses. Lancet 395(10235):1517–1520.

17. Liu Y, et al. (2020) Clinical and biochemical indexes from 2019-nCoV infected patients linked to viral loads and lung injury. Sci China Life Sci 63:364–374.

18. Zitzmann C, Kaderali L (2018) Mathematical analysis of viral replication dynamics and antiviral treatment strategies: From basic models to age-based multi-scale modeling. Front Microbiol 9:1540.

19. Baral S, Antia R, Dixit NM (2019) A dynamical motif comprising the interactions between antigens and CD8 T cells may underlie the outcomes of viral infections. Proc Natl Acad Sci U S A 116(35):17393–17398.

20. Best K, Perelson AS (2018) Mathematical modeling of within-host Zika virus dynamics. Immunol Rev 285(1):81–96.

21. Li K, McCaw JM, Cao P (2020) Modelling within-host macrophage dynamics in influenza virus infection. bioRxiv:083360.

22. Koyama S, Ishii KJ, Coban C, Akira S (2008) Innate immune response to viral infection. Cytokine 43(3):336–41.

23. Qian Z, et al. (2013) Innate immune response of human alveolar type II cells infected with severe acute respiratory syndrome-coronavirus. Am J Respir Cell Mol Biol 48(6):742–748.

24. Pulendran B, Maddur MS (2014) Innate Immune Sensing and Response to Influenza. Influenza Pathogenesis and Control - Volume II, eds Oldstone MBA, Compans RW, pp 23–71.

25. Camp J V., Jonsson CB (2017) A role for neutrophils in viral respiratory disease. Front Immunol 8:550.

26. Brandstadter JD, Yang Y (2011) Natural killer cell responses to viral infection. J Innate Immun 3(3):274–279.

27. Voll RE, et al. (1997) Immunosuppressive effects of apoptotic cells. Nature 390:350–351.

28. Fadok VA, et al. (1998) Macrophages that have ingested apoptotic cells in vitro inhibit proinflammatory cytokine production through autocrine/paracrine mechanisms involving TGF-β, PGE2, and PAF. J Clin Invest 101(4):890–898.

29. Perry AK, Chen G, Zheng D, Tang H, Cheng G (2005) The host type I interferon response to viral and bacterial infections. Cell Res 15:407–422.

30. Tisoncik JR, et al. (2012) Into the Eye of the Cytokine Storm. Microbiol Mol Biol Rev 76(1):16–32.

31. Chigbu DI, Loonawat R, Sehgal M, Patel D, Jain P (2019) Hepatitis C Virus Infection: Host–Virus Interaction and Mechanisms of Viral Persistence. Cells 8(4):376.

32. McLane LM, Abdel-Hakeem MS, Wherry EJ (2019) CD8 T Cell Exhaustion During Chronic Viral Infection and Cancer. Annu Rev Immunol 37:457–95.

33. Johnson PLF, et al. (2011) Vaccination Alters the Balance between Protective Immunity, Exhaustion, Escape, and Death in Chronic Infections. J Virol 85(11):5565–5570.

34. Schmidt ME, Varga SM (2018) The CD8 T cell response to respiratory virus infections. Front Immunol 9:678.

35. Wang Y, Lobigs M, Lee E, Müllbacher A (2003) CD8+ T Cells Mediate Recovery and Immunopathology in West Nile Virus Encephalitis. J Virol 77(24):13323–13334.

36. Kim SE, et al. (2020) Viral kinetics of SARS-CoV-2 in asymptomatic carriers and presymptomatic patients. Int J Infect Dis:in press.

37. Nikolich-Zugich J, et al. (2020) SARS-CoV-2 and COVID-19 in older adults: what we may expect regarding pathogenesis, immune responses, and outcomes. GeroScience:1–10.

38. Honce R, Schultz-Cherry S (2019) Impact of obesity on influenza A virus pathogenesis, immune response, and evolution. Front Immunol 10:1071.

39. Sun B, et al. (2020) Kinetics of SARS-CoV-2 specific IgM and IgG responses in COVID-19 patients. Emerg Microbes Infect 9:940–948.

40. Zheng HY, et al. (2020) Elevated exhaustion levels and reduced functional diversity of T cells in peripheral blood may predict severe progression in COVID-19 patients. Cell Mol Immunol 17:541–543.

41. Lescure FX, et al. (2020) Clinical and virological data of the first cases of COVID-19 in Europe: a case series. Lancet Infect Dis:in press.

42. Yin W, et al. (2020) Structural Basis for the Inhibition of the RNA-Dependent RNA Polymerase from SARS-CoV-2 by Remdesivir. Science:eabc1560.

43. Antunes I, Kassiotis G (2010) Suppression of Innate Immune Pathology by Regulatory T Cells during Influenza A Virus Infection of Immunodeficient Mice. J Virol 84(24):12564–12575.

44. Lin S, Wu H, Wang C, Xiao Z, Xu F (2018) Regulatory T cells and acute lung injury: Cytokines, uncontrolled inflammation, and therapeutic implications. Front Immunol 9:1545.

45. Li S, Gowans EJ, Chougnet C, Plebanski M, Dittmer U (2008) Natural Regulatory T Cells and Persistent Viral Infection. J Virol 82(1):21–30.

46. Lee DCP, et al. (2010) CD25+ Natural Regulatory T Cells Are Critical in Limiting Innate and Adaptive Immunity and Resolving Disease following Respiratory Syncytial Virus Infection. J Virol 84(17):8790–8798.

47. Cox MJ, Loman N, Bogaert D, O’Grady J (2020) Co-infections: potentially lethal and unexplored in COVID-19. The Lancet Microbe:in press.

48. Tay MZ, Poh CM, Rénia L, MacAry PA, Ng LFP (2020) The trinity of COVID-19: immunity, inflammation and intervention. Nat Rev Immunol:in press.

49. Shi Y, et al. (2020) COVID-19 infection: the perspectives on immune responses. Cell Death Differ 27:1451–1454.

50. Prompetchara E, Ketloy C, Palaga T (2020) Immune responses in COVID-19 and potential vaccines: Lessons learned from SARS and MERS epidemic. Asian Pacific J allergy Immunol 38:1–9.

51. Sambaturu N, et al. (2018) Role of genetic heterogeneity in determining the epidemiological severity of H1N1 influenza. PLoS Comput Biol 14(3):e1006069.

52. Kucharski AJ, et al. (2020) Early dynamics of transmission and control of COVID-19: a mathematical modelling study. Lancet Infect Dis 20:553–558.

53. Kaur T, et al. (2020) Anticipating the novel coronavirus disease (COVID-19) pandemic. medRxiv:200057430.

54. Goyal A, Cardozo-Ojeda EF, Schiffer JT (2020) Potency and timing of antiviral therapy as determinants of duration of SARS CoV-2 shedding and intensity of inflammatory response. medRxiv:20061325.

55. Du SQ, Yuan W (2020) Mathematical modeling of interaction between innateand adaptive immune responses in COVIDC19 andimplications for viral pathogenesis. J Med Virol. doi:10.1002/jmv.25866.

56. Perelson AS, Ribeiro RM (2013) Modeling the within-host dynamics of HIV infection. BMC Biol 11:96.

57. Perelson AS (2002) Modelling viral and immune system dynamics. Nat Rev Immunol 2:28–36.

58. Robertson-Tessi M, El-Kareh A, Goriely A (2012) A mathematical model of tumor-immune interactions. J Theor Biol 294:56–73.

59. Padmanabhan P, Garaigorta U, Dixit NM (2014) Emergent properties of the interferon-signalling network may underlie the success of hepatitis C treatment. Nat Commun 5(1):1–9.

60. Pan Y, Zhang D, Yang P, Poon LLM, Wang Q (2020) Viral load of SARS-CoV-2 in clinical samples. Lancet Infect Dis 20(4):P411–412.

61. De Wit E, Van Doremalen N, Falzarano D, Munster VJ (2016) SARS and MERS: Recent insights into emerging coronaviruses. Nat Rev Microbiol 14(8):523–34.

62. Liu L, et al. (2019) Anti-spike IgG causes severe acute lung injury by skewing macrophage responses during acute SARS-CoV infection. JCI insight 4(4):e123158.

63. Kindler E, Thiel V, Weber F (2016) Interaction of SARS and MERS Coronaviruses with the Antiviral Interferon Response. Adv Virus Res 96:219–243.

64. Hu Y, et al. (2017) The Severe Acute Respiratory Syndrome Coronavirus Nucleocapsid Inhibits Type I Interferon Production by Interfering with TRIM25-Mediated RIG-I Ubiquitination. J Virol 91(8). doi:10.1128/jvi.02143-16.

65. Chen X, et al. (2014) SARS coronavirus papain-like protease inhibits the type I interferon signaling pathway through interaction with the STING-TRAF3-TBK1 complex. Protein Cell 5:369–381.

66. Zhou Z, et al. (2020) Heightened innate immune responses in the respiratory tract of COVID-19 patients. Cell Host Microbe:in press.

67. Basak S, Behar M, Hoffmann A (2012) Lessons from mathematically modeling the NF-κB pathway. Immunol Rev 246(1):221–238.

68. Rubio D, et al. (2013) Crosstalk between the type 1 interferon and nuclear factor kappa B pathways confers resistance to a lethal virus infection. Cell Host Microbe 13(6):701–710.

69. Toniati P, et al. (2020) Tocilizumab for the treatment of severe COVID-19 pneumonia with hyperinflammatory syndrome and acute respiratory failure: A single center study of 100 patients in Brescia, Italy. Autoimmun Rev:in press.

