## Supplementary Figures, Methods for "Mechanistic modeling of the SARS-CoV-2 and immune system interplay unravels design principles for diverse clinicopathological outcomes"

**Supplementary Figures and Legends:**

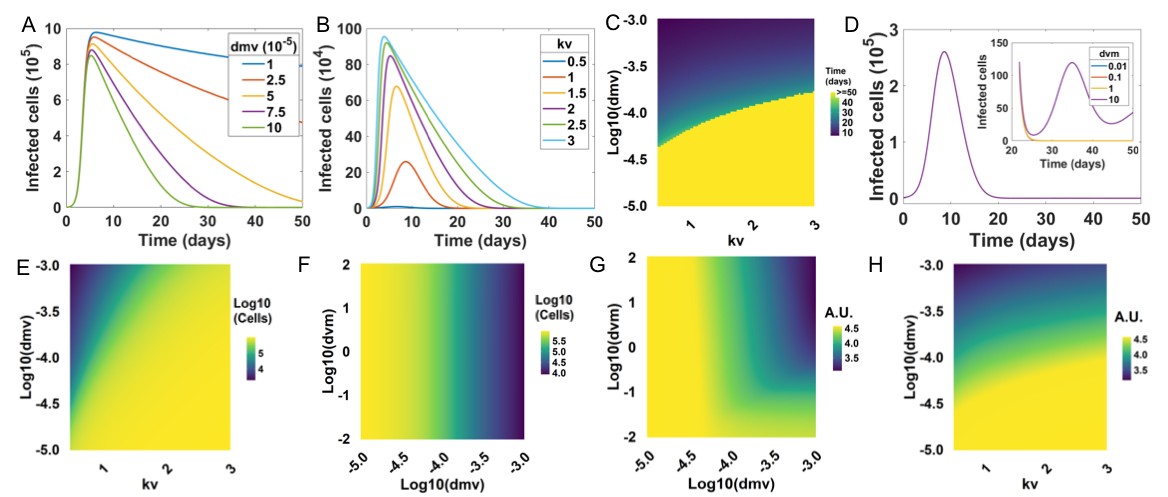

***Fig S1: Sensitivity of viral clearance immunopathology to model parameters. A)*** *Infected cell profiles for varying values of dmv at kv = 2.* ***B)*** *Infected cell profiles for varying values of kv.* ***C)*** *Heatmap showing the change in infection clearance time for multiple combinations of dmv and kv.* ***D)*** *Infected cell profiles corresponding to Fig 1D. Inset shows the zoomed in levels of infected cells after 20 days to highlight the persistence of infection.* ***E)*** *Heatmap showing the dependence of peak of infected cells on kv and dmv.* ***F)*** *Same as E but for dmv and dvm.* ***G)*** *Dependence of peak immunopathology (plotted as log10) on kv and dmv.* ***H)*** *Same as G but for dmv and dvm.*

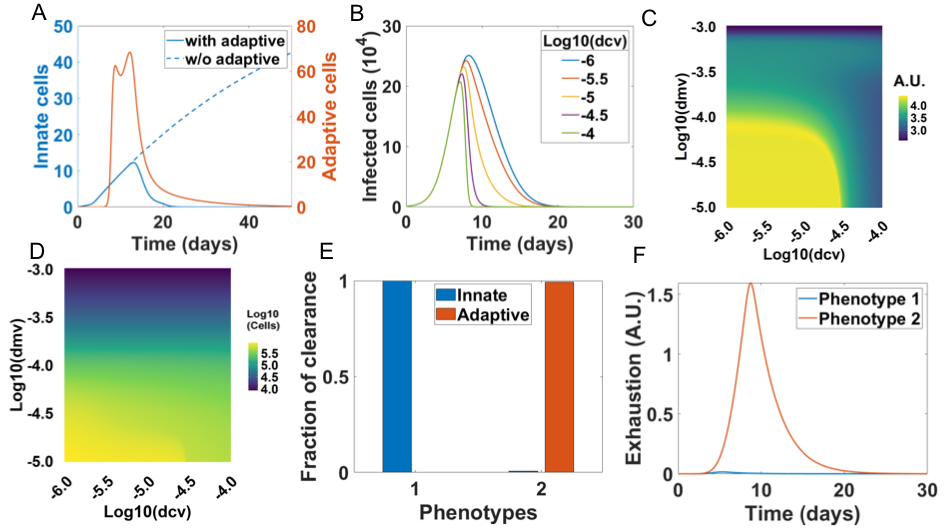

***Fig S2: Sensitivity of viral clearance immunopathology to model parameters. A)*** *Immune cell dynamics corresponding to Fig 2B.* ***B)*** *Infected cell dynamics corresponding to Fig 2C.* ***C)*** *Immunopathology (log10) at 30 days as a function of dmv and dcv.* ***D)*** *Infected cell peak as a function of dmv and dcv.* ***E)*** *Relative contribution of innate (blue) and adaptive (red) immune cells in clearance of infection for the two healthy phenotypes in Fig 2E,F.* ***F)*** *Exhaustion of adaptive cells due to the antigen concentration (Ref: SI eq 9) in the two healthy phenotypes.*

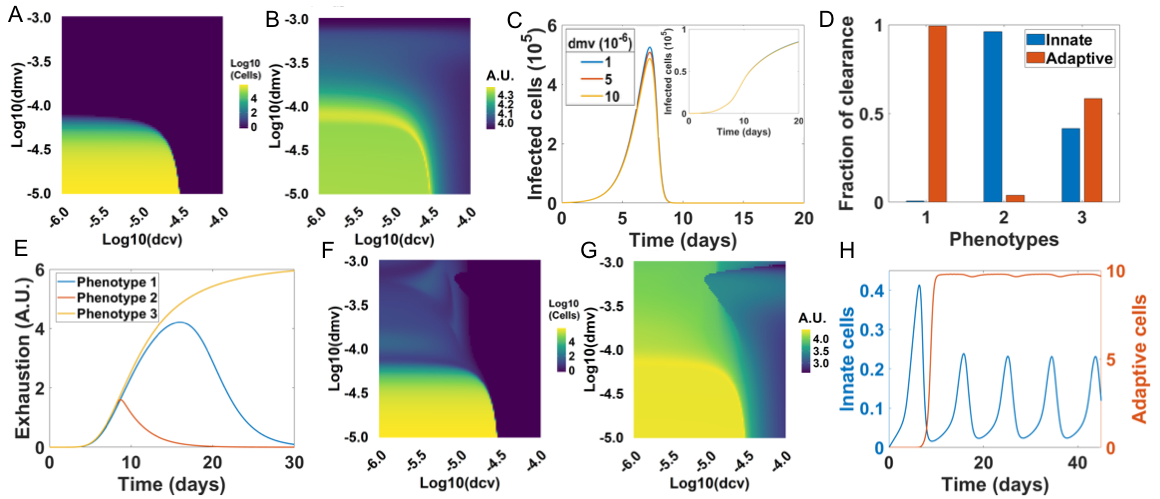

***Fig S3: A)*** *Effect of dmv and dcv on infection load at 30 days at dvm= 0.1.* ***B)*** *Effect of dmv and dcv on immunopathology (log10) at 30 days at dvm = 0.1.* ***C)*** *Infection profiles for varying values of dvm. Inset shows the corresponding immunopathology profiles.* ***D)*** *Relative contribution of innate and adaptive cells in infection clearance the 3 phenotypes in Fig 3A-F.* ***E)*** *Antigen mediated adaptive cell exhaustion dynamics for phenotypes in Fig 3A-F.* ***F)*** *Same as A but at dvm =10.* ***G)*** *Same as B but at dvm = 10.* ***H)*** *Immune cell dynamics for phenotype in Fig 3G-H.*

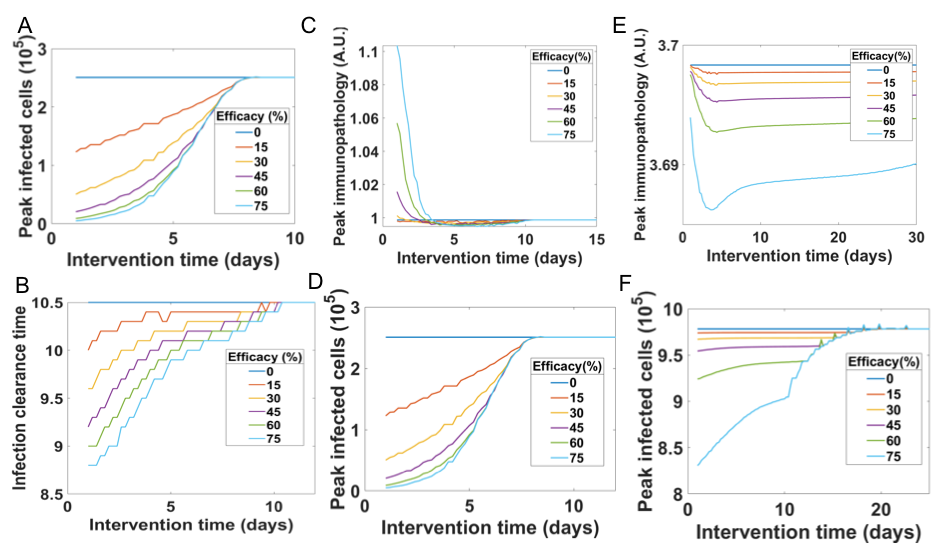

***Fig S4: A)*** *Effect of drug intervention time and efficacy on peak infection levels for severe phenotype 2 (Fig 3C-D).* ***B-D)*** *Effect of drug intervention time and efficacy on clearance time of infection (B), peak immunopathology (C) and peak infection levels (D) corresponding to the severe phenotype 1 (Fig 3A-B).* ***E-F)*** *Same as C-D but corresponding to severe phenotype 3 (Fig 3E-F).*

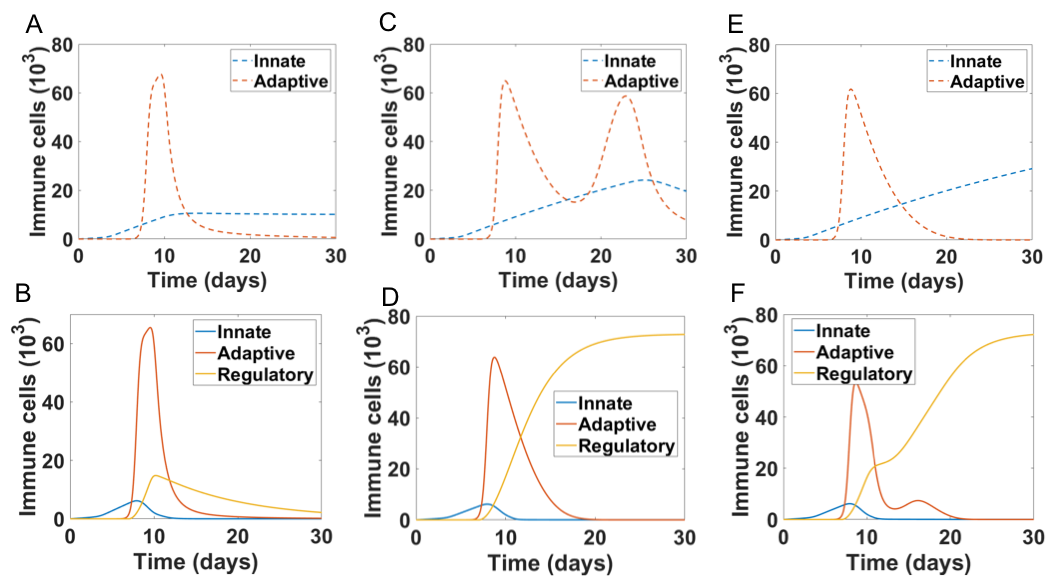

***Fig S5: A-B)*** *Immune cell profiles for 5B without (A) and with (B) regulatory cells.* ***C-D)*** *: Immune cell profiles for 5C without (C) and with (D) regulatory cells.* ***E-F)*** *: Immune cell profiles for 5D without (E) and with (F) regulatory cells.*

**Supplementary Methods:**

The mathematical model was simulated using the ode45 ODE solver in MATLAB R2018b. The heatmaps were plotted using ggplot2 in R 3.6. All other graphs were generated using MATLAB. Default values for the parameters have been used for all the simulations (given in Supplementary table 2) unless mentioned otherwise.

**1. Interactions between the innate immune system and the SARS-CoV-2 infected cells.**

The interactions between the innate and the infected cell populations can be modelled via the following ordinary differential equations:

$$\frac{dV}{dt}=H\left( V-1 \right)*\left( kv*V*\left( 1-\frac{V}{V_{max}} \right)-dmv*M*V \right)-\left( 1-H\left( V-1 \right) \right)*V (1)$$

$$\frac{dM}{dt}=kvm*\frac{V}{qvm+V}+kmm*\frac{M^{n}}{qmm^{n}+M^{n}}-dvm*M*\frac{qvm2}{qvm2+V}-dm*M (2)$$

$$\frac{dP}{dt}=kmp*M+kvp*V-dp*P (3)$$

where, *V* is the number of virally infected cells, *M* is the number of innate immune cells and *P* is the cytokine mediated immunopathology.

The dynamics of the variable V indirectly include the dynamics of viral load as well. H(V-1) is the Heaviside function, defined as 0 when V<1 and 1 when V>1. This function describes the condition for infection clearance. The SARS-CoV-2 virally infected cells can be assumed to grow at a rate of *kv* in a logistic fashion to reach a carrying capacity, V_max_, a measure of the number of susceptible cells in the system. The infected cells can be cleared by the innate immune cells (M) at a rate of dmv. Furthermore, a first-order decay of infection replaces the other dynamic terms above upon clearance, i.e., V<1.

On the other hand, the dynamics of innate immune cells (M) can be modelled as a “biphasic process”. These cells can either be recruited at a maximal rate of kvm by the virally infected cells through the secretion of cytokines and chemokines or proliferate at a maximal rate of kmm due to the stimulation by other innate immune cells or due to pro-inflammatory cytokines and chemokines. Furthermore, innate immune cells have the ability to self-inhibit when there is a very low load of the infected cells in the system (resolution of innate inflammation). This can be captured by a negative Hills’ functions with a maximal rate of dvm. The innate immune cells can also be cleared at a rate of dm following first-order kinetics.

The level of immunopathology in the system (P) is by the contributions from infected cells (V) and innate immune cells (M). The virally infected cells contribute to the immunopathology at a rate of kvp while the innate immune cells contribute to the immunopathology at a rate of kmp. It is of importance to note that we assume that the rate kvp << kmp, as innate immune cells such as the macrophages, monocytes, neutrophils, natural killer cells, etc. are likely more potent at creating a pro-inflammatory environment than the virally infected cells. The system can recover from the immunopathology at a rate of dp again approximated by first order decay kinetics.

**2. Interactions between the adaptive immune system and the SARS-CoV-2 infected cells.**

To incorporate the interactions between the adaptive immune cells (C) with the virally infected cells (V), we modified the above set of equations in the following manner:

$$\frac{dV}{dt}=H\left( V-1 \right)*\left( kv*V*\left( 1-\frac{V}{V_{max}} \right)-dmv*M*V\boldsymbol{-dcv*C*V} \right)-\left( 1-H\left( V-1 \right) \right)*V (4)$$

$$\frac{dM}{dt}=kvm*\frac{V}{qvm+V}+kmm*\frac{M^{n}}{qmm^{n}+M^{n}}-dvm*M*\frac{qvm2}{qvm2+V}-dm*M (5)$$

$$\frac{dC}{dt}=lamb1*H\left( t-t_{C} \right)+ kvc*\frac{C*V}{qvc+V}*\left( 1-\frac{C}{C_{max}} \right)-dqc*\frac{C*Q^{n}}{Q^{n}+qqc^{n}}-dc*C (6)$$

$$\frac{dQ}{dt}=kvq*\frac{V}{qvq+V}-dq*Q (7)$$

$$\frac{dP}{dt}=kvp*V+kmp*M\boldsymbol{+kcp*C}-dp*P (8)$$

In this scenario the infected cells can be cleared via another mechanism; killing by adaptive cells at a rate of dcv. Furthermore, dynamics of antigen load has been included such that antigen load (Q) accumulates at a rate of kvq with a sigmoidal dependence on the infected cell number. The antigen load decays following a first order rate kinetics at a rate of dq.

The adaptive cells themselves infiltrate into the site of infection at a constant rate of lamb1 after a time of tc days. This is in accordance with the fact that the adaptive immune response sets in after a few days in contrast to the innate immune system that is triggered almost instantaneously. The adaptive cells initially proliferate rapidly at a rate of kvc in the presence of virally infected cells (V) mediated by the antigen presenting cells. The proliferation also has a logistic component to it accounting for the clonal selection of adaptive immune cells and for the fact that the site of infection can accommodate at most a fixed number of cells. The adaptive cells can get exhausted at a maximal rate of dqc depending on the antigen load (Q) in the system. Specifically, we define the antigen-mediated exhaustion as:

$$Exhaustion \left( at time t \right)=dqc*\frac{Q^{n}}{Q^{n}+qqc^{n}}---(9)$$

Where Q is the instantaneous antigen accumulated at time t. Furthermore, the adaptive cells get killed or removed from the site of the infection at a rate of dc.

The adaptive immune cells contribute to the immunopathology at a rate of kcp. Again, we have kvp < kcp < kmp as adaptive cells are known to cause immunopathology, they do not contribute to it as much as the innate immune cells. Along with that there are a few reports the suggest adaptive cells might even secrete IL-10, a well-known anti-inflammatory cytokine along with pro-inflammatory cytokines, making their net contribution to immunopathology lesser than the innate cells. Adaptive cells however are assumed to have a greater potency to cause immunopathology than the virally infected cells.

**3. Including the effect of regulatory cells**

To incorporate the effects of the regulatory cells (R) (more specifically the regulatory T cells) we modified the set of equations to the following:

$$\frac{dV}{dt}=H\left( V-1 \right)*\left( kv*V*\left( 1-\frac{V}{V_{max}} \right)-dmv*M*V-dcv*C*V \right)-\left( 1-H\left( V-1 \right) \right)*V---(10)$$

$$\frac{dM}{dt}=kvm*\frac{V}{qvm+V}+kmm*\frac{M^{n}}{qmm^{n}+M^{n}}-dvm*M*\frac{qvm2}{qvm2+V}-\boldsymbol{drm*M*R-}dm*M---(11)$$

$$\frac{dC}{dt}=lamb1*H\left( t-t_{C} \right)+ kvc*\frac{C*V}{qvc+V}*\left( 1-\frac{C}{Cmax} \right)-dqc*\frac{C*Q^{n}}{Q^{n}+qqc^{n}}-\boldsymbol{drc*C*R}-dc*C---(12)$$

$$\frac{dQ}{dt}=kvq*\frac{V}{qvq+V}-dq*Q---(13)$$

$$\frac{dR}{dt}=lamb2*H\left( t-t_{R} \right)+ kvr*\frac{R^{0.5}*V}{qvr+V}*\left( 1-\frac{R}{Rmax} \right)-dr*R---(14)$$

$$\frac{dP}{dt}=kvp*V+kmp*M+\boldsymbol{kcp*C}-dp*P---(15)$$

Similar to the adaptive immune cells we assume that the regulatory T cells would infiltrate at time tR with a rate lamb2 and proliferate with a rate kvr to a carrying capacity of Rmax. However, the regulatory cells are known not to proliferate as rapidly as the clonal expansion of adaptive cells, hence we have a sub-linear dependence on the regulatory cells (R). Here we assume the exhaustion of the regulatory cells is low at the time scales of the model and hence it is not considered. Furthermore these cells can die or be cleared at a rate of dr from the site of the infection.

**4. Modelling the effect of antiviral drugs:**

Antiviral drugs reduce the virulence of infection, represented by the model parameter kv. To model this effect, we replace kv in the previous set of equations with kv’ such that:

$$k_{v}^{'}=k_{v}*\left( H\left( tau-t \right)+H\left( t-tau \right)*\left( 1-e*0.01 \right) \right)---(16)$$

Here, tau is the time of administration of the drug and e is the efficacy of the drug, i.e., the percentage reduction in the growth rate of the infection due to the drug.
